# Dealing with lipid effects and lipid-extraction biases in δ^13^C and δ^15^N isotopic studies: a solution based on 28 marine invertebrate, fish and mammal species

**DOI:** 10.1101/2023.10.25.563823

**Authors:** Jean-François Ouellet, Jory Cabrol, Ève Rioux, Xavier Bordeleau, Véronique Lesage

## Abstract

Lipids are naturally depleted in ^13^C isotope in relation to its C sources, causing a bias in δ^13^C in bulk samples that varies with lipid content. Failure to take this issue into account results in inaccurate conclusions in food web and habitat use studies. Two approaches to resolve this issue are 1) to extract lipids from samples prior to measurement, a resource-intensive process that also can alter δ^15^N or 2) estimating a lipid-free δ^13^C using one of several equations that differ in levels of sophistication and generalization across taxa. Here δ^13^C and δ^15^N were measured on bulk and lipid-extracted muscle samples of a dataset of over 2000 specimens of 28 species of marine invertebrates, fishes and mammals. Our objectives were to 1) compare the effect of lipid extraction on δ^13^C and δ^15^N across taxa; 2) compare the performance of five normalization models, overall and on subsets of species; 3) propose a model to revert lipid-extracted δ^15^N back to their bulk values; and 4) identify the most suitable approach for dealing with lipid biases in isotopic ratios. Extraction caused an uneven enrichment in δ^13^C and δ^15^N across species. Model taxonomic specificity increased estimate accuracy in both isotopes. Models from Logan et al. (2008) and McConnaughey and McRoy (1979) performed better than the other models tested. δ^15^N_bulk_ could be reliably estimated based on δ^15^N_lipid-extracted_ using a linear model. This study provides a way forward for obtaining reliable δ^13^C and δ^15^N values in muscle tissue without the costs of duplicate analyses and represents a major step toward the harmonization of datasets collected under the two different approaches.

## 1. INTRODUCTION

Stable isotopes of various chemical elements that are present in the environment vary in abundance with landscape and ecosystem features. Some isotopes such as the lighter ^12^C and ^14^N are naturally abundant, and generally metabolized in animal tissues and excreted preferentially over their heavier form (e.g., ^13^C and ^15^N; Peterson & Fry 1987, Hobson et al. 1996). As a result, heavier isotopes tend to accumulate more in animal tissues relative to the lighter ones making them useful tracers of carbon sources, trophic position, and thereby of habitat use and movement patterns across isoscapes (Hobson 1999, Post 2002, Perrin et al. 2014).

Despite their broad utilization in trophic ecology studies, an important confounding factor of stable isotope analyses has been variability in the lipid content of samples, naturally depleted in ^13^C relative to other biochemical compounds (DeNiro & Epstein 1977). The presence and variability of lipids within and among species may lead to misinterpretation of changes in diet or habitat use. While the main stance during the early developments of the method has been to ignore the issue, consensus quickly built about the need to assess potential effects of lipids on δ^13^C values and if significant, to account for this bias (Hobson et al. 1996, Post et al. 2007, Choy et al. 2016). For about 15 years (e.g., mid-1990s to the mid-to late 2000’s), lipid extraction prior to isotope analysis was the most widespread approach. This procedure was based on the assumption that an absence of nitrogenous components in chemical solvents would prevent alteration of δ^15^N values. However, this assumption was refuted by empirical assessments, with multiple studies reporting significant and uneven discrimination, resulting in ^15^N enrichment, across species following lipid extraction (Choy et al. 2016, Clark et al. 2019, Lerner & Hunt 2022, but see Cloyed et al. 2020). An alternative to lipid extraction has been to analyze carbon and nitrogen isotopes using separate aliquots (Logan et al. 2008, Lesage et al. 2010). The consequent doubling of analytical costs and time increased the interest for lipid normalization models for δ^13^C values, leading to numerous proposed approaches with various levels of sophistication and generalization (McConnaughey & McRoy 1979, Fry et al. 2003, Logan et al. 2008, Lesage et al. 2010, Clark et al. 2019, Fischer-Rush et al. 2021). Empirical comparison of models showed various levels of performance across taxonomic groups and tissues (Logan et al. 2008), emphasizing the importance of model validation and specificity. Variability in the ways investigators handle the issue of ^13^C depletion in lipids may hinder data comparisons between studies. As a result, caution and validation are needed when selecting an adequate model for the taxa or tissue under study, including coefficient values. But thus far, little guidance exists for investigators in making that choice. Also, equivalent models for reverting lipid-extracted δ^15^N values back to their bulk un-extracted values have been published (Logan et al. 2008, Lesage et al. 2010, Groß et al. 2021, Lerner & Hunt 2022), making older datasets, where lipid-extraction was the norm, suitable for long-term monitoring of environmental or trophic change.

Here, we take advantage of a dataset of over 2,000 specimens from 28 marine species including invertebrates, bony and cartilaginous fishes, and marine mammals for which both lipid-extracted and bulk δ^13^C and δ^15^N values have been determined to improve our understanding of these questions. Specifically, we 1) compared the effect of chemically extracting lipids on δ^13^C and δ^15^N across taxa; 2) examined the relative performance of five popular mathematical normalization models for estimating δ^13^C_lipid-free_, both overall and for subsets of resembling species; 3) proposed a helpful model to restore old datasets where lipid extraction was applied systematically to all samples, and revert lipid-extracted δ^15^N values back to their bulk values; 4) identified the most suitable approach for dealing with lipid effects on isotopic values, including the best models and parameters to use for a given dataset or particular taxa. This study represents a major step toward the harmonization of datasets collected under different approaches for long-term studies.

## 2. MATERIAL AND METHODS

### 2.1 Sample collection

Our sample consisted of 2,354 muscle samples from 28 marine species of invertebrates, bony and cartilaginous fishes, and marine mammals (Table 1), collected in 1997—2021 in the Estuary and Gulf of St. Lawrence, Canada. Most of the fish and invertebrate specimens were obtained through an annual trawling mission (Bourdages et al. 2019). Additional samples were obtained from various scientific missions and sampling programs, commercial fisheries using fish traps or by opportunistically netting capelin (*Mallotus villosus*), krills, and sand lance (*Ammodytes* sp.) from small vessels. Beluga (*Delphinapterus leucas*) samples were obtained post-mortem from well-preserved to moderately-decomposed beach-cast carcasses (freshness code 2 or 3, Geraci & Lounsbury 1993, Lesage 2014), whereas grey seal (*Halichoerus grypus*) samples were obtained from scientific or commercial harvests conducted in the Magdalen Islands (Gulf of St. Lawrence) between 2010 and 2021. All samples were frozen upon collection in air-tight plastic bags and stored at −20°C.

**Table 1.**
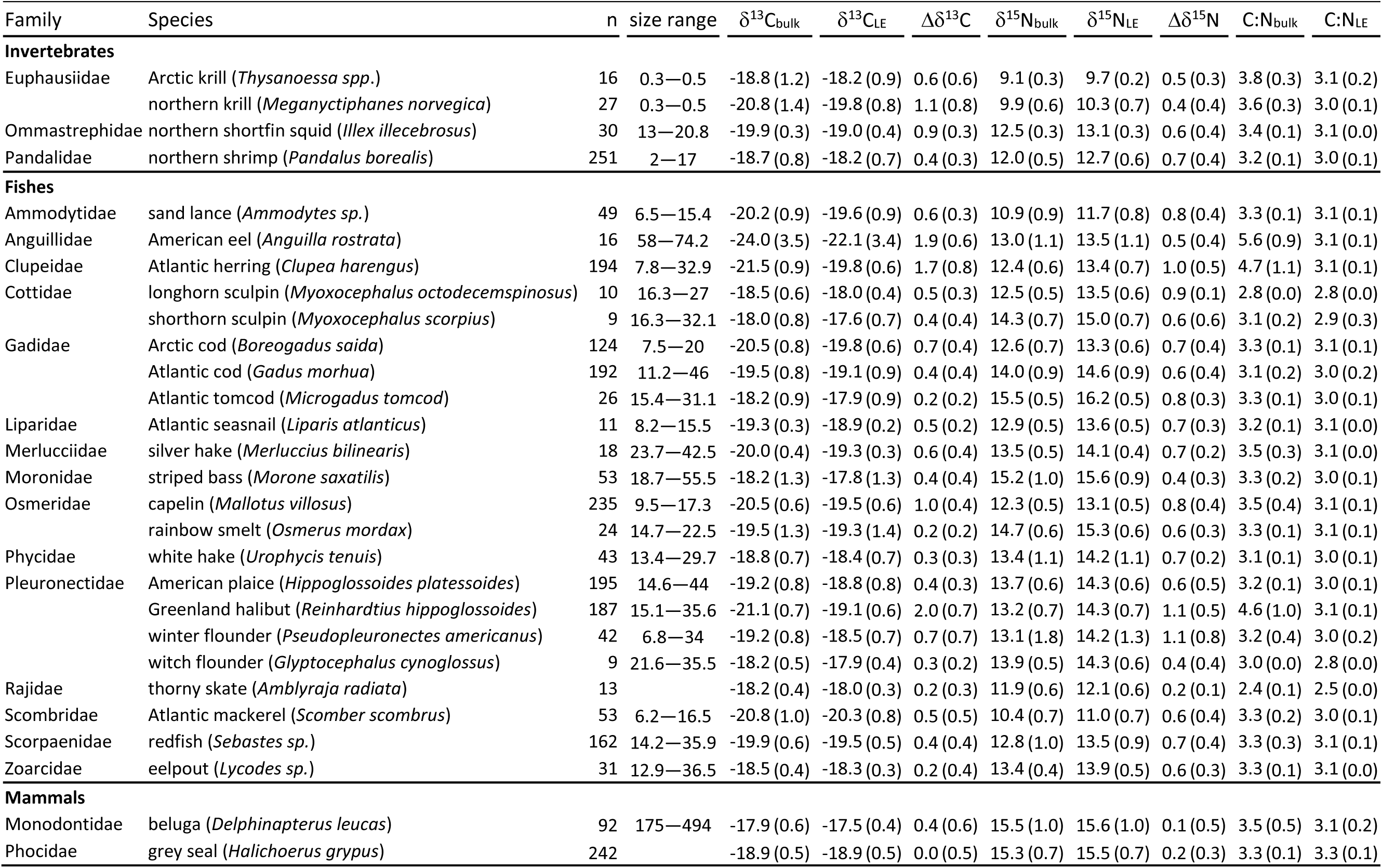
Sample size (*n*), size range (cm) of specimens and mean (*SD*) values (‰) of carbon and nitrogen stable isotopes ratios (δ^13^C, δ^15^N) and discrimination (Δδ^13^C, Δδ^15^N) and C:N values measured in bulk and lipid-extracted (LE) muscle samples from species under study.

Muscle samples were excised dorso-laterally from fishes (∼1.5 g), grey seal (∼1.5 g) and beluga (∼500 cm^3^), while the whole dorsal muscle without the carapace was taken from krills and northern shrimp (*Pandalus borealis*), and the mantle was taken from northern shortfin squid (*Illex illecebrosus*). Samples were minced, freeze-dried for 48 h, and ground to a fine powder with a mortar and pestle (Wig-L-Bug, Crescent Dental Mfg.). Samples were then divided in two aliquots: one aliquot received no further treatment prior to isotope analysis (bulk); the second aliquot was lipid-extracted.

Lipids were extracted using a 2:1 (v/v) chloroform and methanol mixture following the procedure described by Folch et al. (1957) and outlined in Lesage et al. (2010). Briefly, ∼0.2 g of powdered muscle was mixed with 10 ml of the chloroform and methanol mixture, sonicated for 15 min, and stored overnight at 4°C with gentle shaking. The sample was then centrifuged for 10 min at 1,500 rpm and the supernatant discarded. The lipid extraction process was repeated twice, at the exception that the mixture was put on the agitator plate for one hour instead of overnight. After extraction, samples were desiccated by evaporation and dried overnight at 60°C.

Subsamples of 0.350 to 0.500 mg of homogenized muscle tissue were precisely weighted and encapsulated in tin capsules. δ^13^C and δ^15^N values, along with the percentage of carbon (%C) and nitrogen (%N) content, were measured in bulk and lipid-extracted aliquots of each sample using a continuous-flow stable-isotope mass spectrometer coupled to a Carlo Erba elemental analyzer (CHNS-O EA1108). Isotopes ratios are expressed in δ notation in parts per thousand (‰) difference from a standard (Vienna Pee Dee Belemnite limestone and atmospheric N for ^13^C and ^15^N, respectively) and calculated as follows:

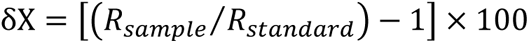

where X is ^13^C or ^15^N and R is the corresponding ^13^C/^12^C or ^15^N/^14^N value. C:N was calculated as the ratio of %C on %N.

The accuracy of isotopic analyses was assessed using commercially certified material (acetanilide or nicotinamide), whereas precision was assessed by replicating measurements every ten samples in the series. The analytical error was ± 0.1‰ for δ^13^C and ± 0.3 for δ^15^N based on laboratory standards, whereas the repeatability for replicates was 0.2‰ for δ^13^C, 0.3‰ for δ^15^N, 0.92% for %C, and 0.31% for %N (*n* = 471).

### 2.2 Statistical analyses

Paired *t*-tests were used to examine, for each species, the effect of lipid extraction on δ^13^C, δ^15^N and C:N. The difference between lipid-extracted and bulk aliquots was expressed as Δδ^13^C (δ^13^C_lipid-extracted_ −δ^13^C_bulk_) and Δδ^15^N (δ^15^N_lipid-extracted_ - δ^15^N_bulk_). ^13^C discrimination is known to be related to C:N_bulk_ in a log-linear relationship (Logan et al. 2008, Lesage et al. 2010, Fischer-Rush et al. 2021, Lerner & Hunt 2022), and this relationship was tested here.

The potential for lipid normalization of δ^13^C values using δ^13^C_bulk_ was estimated with five models that have been commonly used in the literature over the past decade. These models varied both in structure (linear and non-linear models) and in the set of constant parameters they included. The MM model (McConnaughey & McRoy 1979) is a non-linear model developed using the whole body of various marine vertebrates and invertebrates:

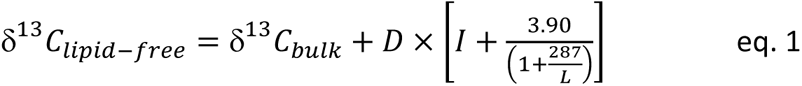

where,

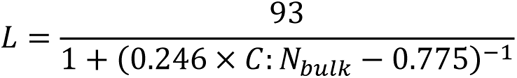

and where I is a constant (equal to −0.207), *L* is the lipid content, and *D* is the isotopic difference between pure protein and pure lipids (assumed to be 6‰, McConnaughey & McRoy 1979), respectively.

The Fry model (Fry 2002), customarily referred to as the mass-balance approach, was developed using muscle of freshwater fish species:

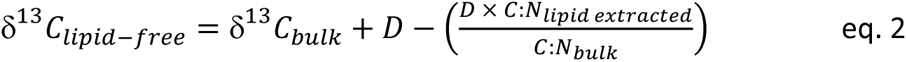

where D is again the isotopic difference between pure protein and pure lipids.

The Post model (Post et al. 2007; Eq. 3), is a linear model developed using aquatic species (muscle or whole organisms):

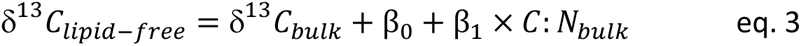

where β_0_ = −3.32 and β_1_ = 0.99.

The Logan model (Logan et al. 2008) is a log-linear relationship developed using freshwater and marine species, and multiple tissues:

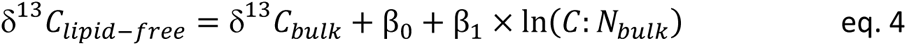

where β_0_ = −4.763 and β_1_ = 4.401 for fish muscle tissue.

Finally, the Lesage model (Lesage et al. 2010) is a linear model developed for cetacean skin. It is the sole model tested that did not take C:N into account:

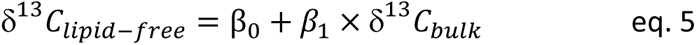

where β_0_ = −4.334 and β_1_ = −0.331.

In contrast with carbon stable isotope for which we seek to estimate δ^13^C_lipid-free_ values, for nitrogen isotopes, it is generally δ^15^N_bulk_ that we seek to estimate and restore from lipid-extraction biases.Because δ^15^N_bulk_ is expected to be linearly related to δ^15^N_lipid-extracted_ (Logan et al. 2008, Lesage et al. 2010, Groß et al. 2021, Lerner & Hunt 2022), this relationship was tested here (see Results) to accurately retro-estimate δ^15^N_bulk_ from δ^15^N_lipid-extracted_.

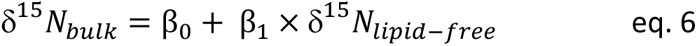

### 2.3 Model performance

Model performance was assessed by linearly regressing predicted values against observed values (δ^13^C_lipid-extracted_ or δ^15^N_bulk_) and examining the mean absolute error and accuracy in predictions, calculated as the relative proportion of predicted values falling within the mean measurement error (0.2‰ and 0.3‰ for δ^13^C and δ^15^N, respectively) of observed values. Also, the slopes of these relationships were compared with a target value of 1.00. The coefficient of determination (r^2^) and Akaike information criterion (AIC) were calculated for within-group comparison (see below) of carbon normalization models.

### 2.4 Model specificity

Precision of estimates and model fit are expected to improve with sample size but also with data specificity (i.e., specific tissue and species, Logan et al. 2008, Lesage et al. 2010, Cloyed et al. 2020). In an attempt to fulfill these conflicting conditions, a trade-off was reached by dividing the sample in clusters of resembling species. Here, species resemblance was defined as a similarity in the species response to lipid extraction rather than phylogenetic kinship. Normalization models were therefore run according to 4 scenarios that differed in sample size and data specificity. The comparison of model performance across scenarios allowed to check if doing so actually improved models fit. The global approach scenario aimed at maximizing sample size: all specimens were combined and the models were run on a single cluster of 2,354 specimens. In the cluster-based scenario, the models were run on smaller clusters of species that were defined using mixture models (R package flexMix, Leisch 2004, Grün & Leisch 2007, 2008). Mixture models are a convenient tool for distributing items in an optimized number of clusters on the basis of the similarities in their relationship with an independent variable (Hamel et al. 2017). Here, the clusterization was based on the relationship of δ^13^C_bulk_ against C:N_bulk_ for δ^13^C, and that of δ^15^N_bulk_ against δ^15^N_lipid-extracted_ for δ^15^N. The optimal number of clusters (k) was determined using the stepFlexMix function (R package flexMix) and Bayesian information criterion (BIC) after repeated trials with candidate values of k ranging from 1 to 28, the maximum number of species. In the species-specific scenario, each species was individually considered as a cluster, maximizing model specificity. In the original coefficients scenario, the models were run on clusters of species like in the cluster-based scenario, but using the originally published coefficient values (presented above with each equation), reducing thereby model specificity to our particular dataset. In the global approach, cluster-based and species-specific scenarios, least-squares estimates of coefficients were obtained for each combination of model and cluster with function nls2 (R package nls2, Grothendieck 2022) for predicting δ^13^C_lipid-free_, and with the function refit (R package flexMix) for predicting δ^15^N_bulk_. The performance of the various models and scenarios for grouping species or estimating coefficients was evaluated by comparing results with a model where all specimens were included into a single linear regression (predicted vs observed value). All statistical analyses were performed using R software version 4.2.2 (R Core Team 2021). Homogeneity of variance and normality assumptions were tested for each statistical analysis where relevant.

## 3. RESULTS

### 3.1 Effect of lipid extraction on isotopes and C:N ratios

Lipid extraction caused a significant enrichment in both δ^13^C and δ^15^N values for all species (paired *t*-tests: *t* >2.2, *p* <0.05 for each species) except for δ^13^C in grey seal (*Halichoerus grypus*) (*t* = −0.18, *p* = 0.86). Mean discrimination varied across species from 0.0 to 2.0‰ for δ^13^C and from 0.1 to 1.1‰ for δ^15^N (Table 1; Fig. 1a, b) and exceeded measurement errors in all cases with the exception of grey seal and thorny skate (*Amblyraja radiata*) for δ^13^C (Fig. 1a), and grey seal, thorny skate and beluga (*Delphinapterus leucas*) for δ^15^N (Fig. 1b). Species showing the largest δ^13^C enrichment were predictably those with the highest C:N_bulk_ values, as shown by the non-linear relationship between these variables (Fig. 1c). Such a relationship with C:N_bulk_ was non-existent for δ^15^N (F_1,26_ = 1.5, *p* = 0.23, not shown).

**Figure 1.**
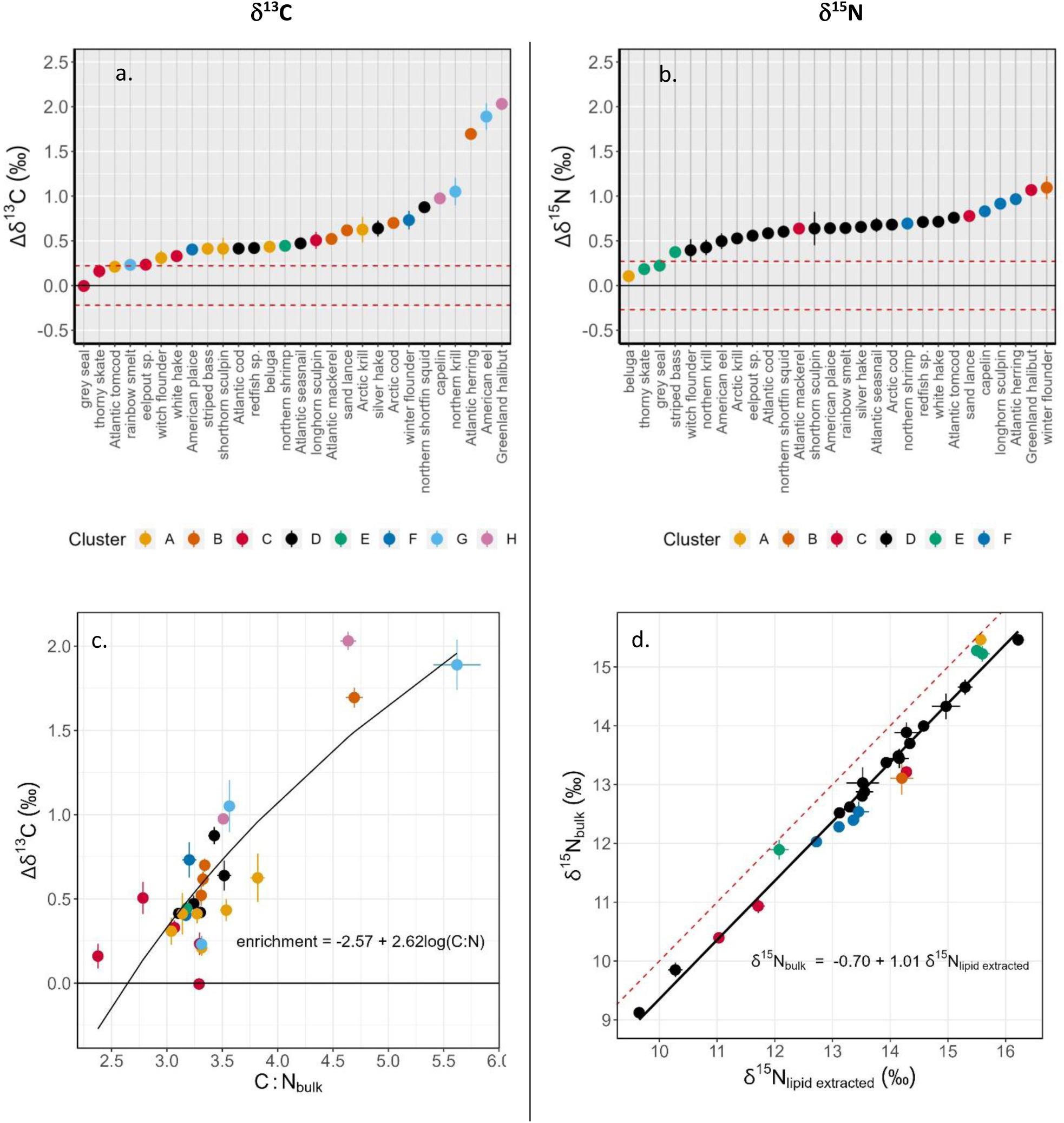
Mean ± *SE* discrimination following lipid extraction (lipid-extracted - bulk) for (a) δ^13^C and (b) δ^15^N, in 28 marine species. Species are sorted by mean value. Horizontal dashed lines indicate the upper 95% confidence limit of intra-individual difference between replicates, above and below 0 ‰. (c) Relationship between mean (± *SE*) δ^13^C discrimination and C:N_bulk_. (d) Relationship between mean ± *SE* bulk and lipid-extracted δ^15^N for 28 marine species. Red dashed line indicates a relationship with slope 1 and intercept 0. See Table 1 for sample sizes.

However, our data showed, as expected based on published studies (e.g., Logan et al. 2008, Lesage et al. 2010, Groß et al. 2021, Lerner & Hunt 2022), a linear relationship between δ^15^N_bulk_ and δ^15^N_lipid-extracted_ (Fig. 1d).

### 3.2 Identifying groups of resembling species

Based on stepFlexMix function and lowest Bayesian information criterion (BIC), the optimal number of clusters of resembling species within our dataset was eight for δ^13^C and six for δ^15^N. Posterior probabilities (i.e., the probability for a species to be adequately assigned to a cluster, indicative of the robustness of the clusterization output) were high for both isotopes, with means and medians of 0.95 and 1.00 for the classification based on carbon (C-classification), and 0.92 and 0.99 for the classification based on nitrogen (N-classification). The relationships used to discriminate the clusters (δ^13^C_bulk_ vs C:N_bulk_ and δ^15^N_bulk_ vs δ^15^N_lipid-extracted_) were generally negative for the C-classification with the exception of cluster E, which was composed of northern shrimp (*Pandalus borealis*) only, and positive for the N-classification (Fig. 2). Clusters differed in species composition between the C- and N-classifications, and in both cases, showed little apparent taxonomic correlation within clusters: species belonging to the same family were seldom clustered together (Fig. 3).

**Figure 2.**
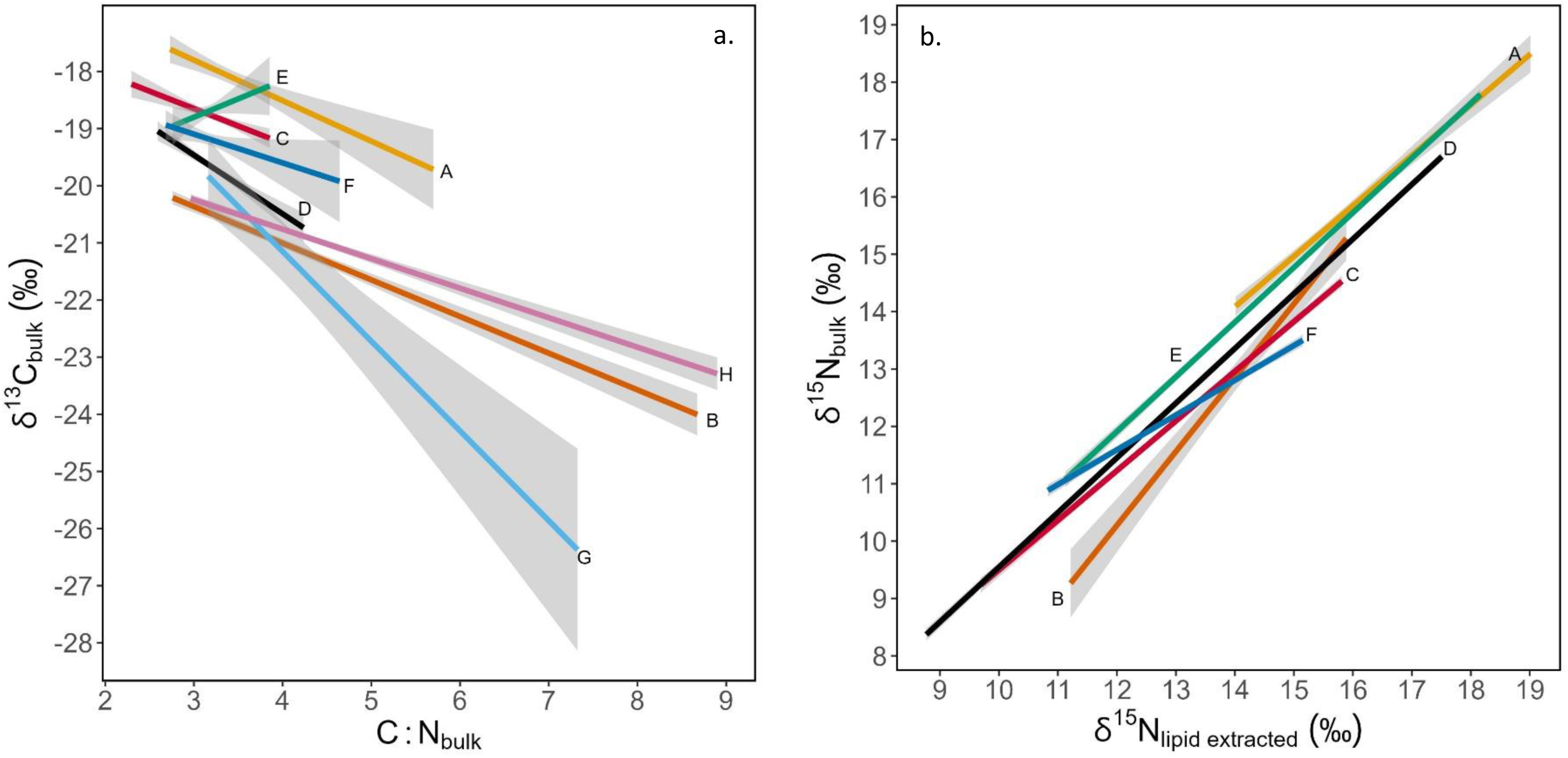
Linear relationships (± *SE*) forming the basis for the clusterization procedure in a) the C-classification (δ^13^C_bulk_ *vs* C:N_bulk_), and b) the N-classification (δ^15^N_bulk_ *vs* δ^15^N_lipid extracted_), for each cluster of species. Clusterization was performed using stepFlexmix function (R package flexMix, Leisch 2004, Grün & Leisch 2007, 2008). Labels indicate clusters names. Cluster composition differs between both classifications.

**Figure 3.**
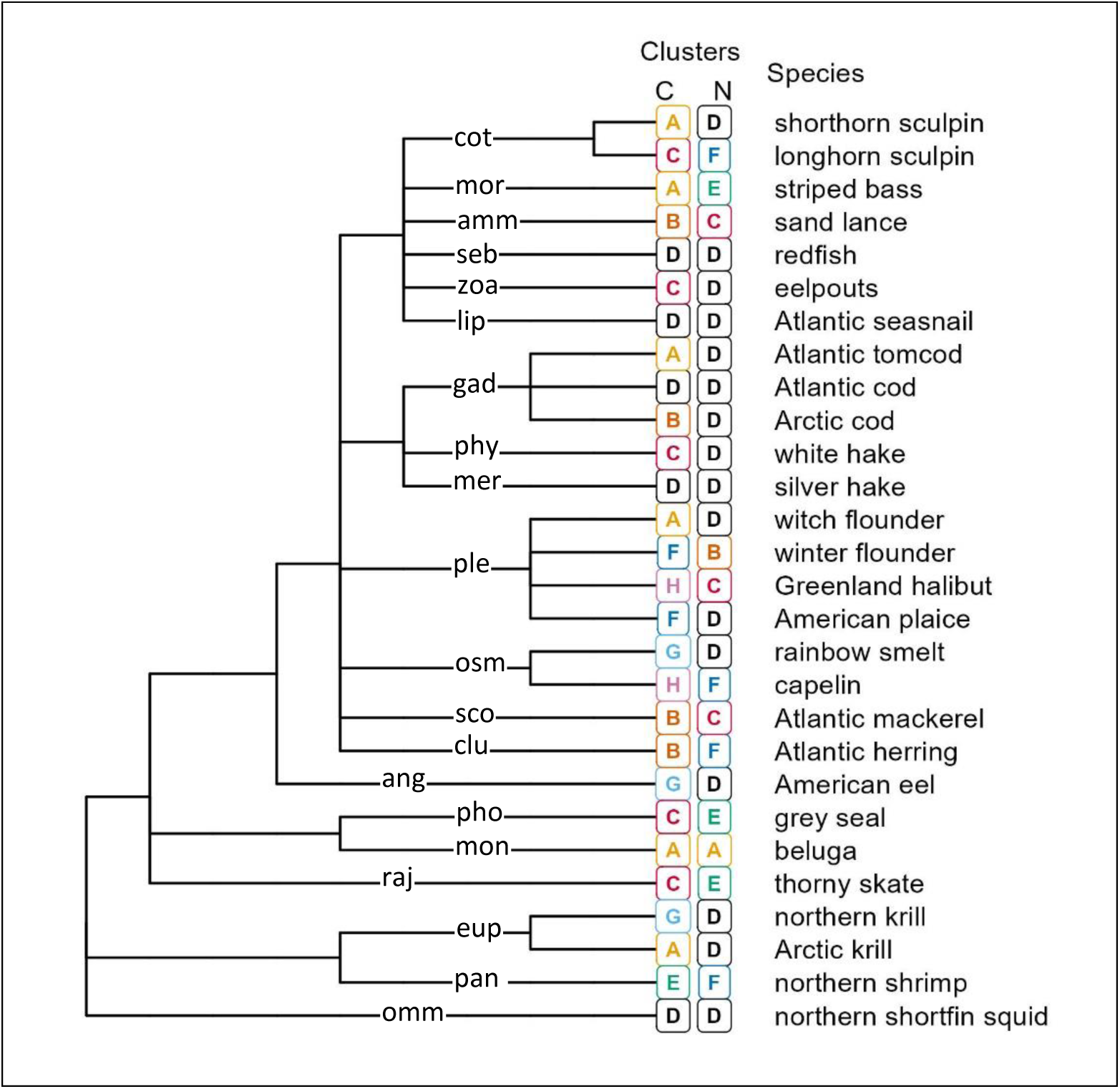
Dendrogram showing the phylogeny of the 28 species in the sample and the clusters to which the species belong. C and N indicate the classifications based on carbon and nitrogen, respectively, and produced with mixture models using stepFlexmix function (R package flexMix, Leisch 2004, Grün & Leisch 2007, 2008). Families are abbreviated with first three letters, full names are as follows: ammodytidae, anguillidae, clupeidae, cottidae, euphausiidae, gadidae, liparidae, merlucciidae, monodontidae, moronidae, ommastrephidae, osmeridae, pandalidae, phocidae, phycidae, pleuronectidae, rajidae, scombridae, sebastidae, zoarcidae. Label placement for families along x axis are arbitrary. Phylogenetic data were extracted from World Register of Marine Species (WoRMS)

The comparison of model performances across all scenarios, for the two isotopes, showed that regardless of the lipid-normalization model selected, model fit was invariably the best in the species-specific scenario (Table 2). Model performances were consistently the poorest in the global approach scenario, with a 0.1‰ average increase in mean absolute error and an average loss of relative accuracy of 11% compared to the species-specific scenario. The cluster-based scenario was intermediate between global approach and species-specific scenarios but showed a smaller deviance from species-specific than the global approach scenario (Table 2), with an average increase in the mean absolute error of 0.03‰ and average decrease in relative accuracy of 4%.

**Table 2.** Comparison of models performance according to four scenarios that differ in level of data specificity (global approach, cluster-based, species-specific, original-coefficients scenarios). mae : mean absolute error (‰), diff_mae_ : difference between the mean absolute error in a scenario and that of the best scenario within the same model (which was invariably the species-specific scenario), acc.: relative accuracy (%, the relative proportion of predicted values falling within the mean measurement error), diff_acc_ : difference between the relative accuracy in a scenario and that of the best scenario within the same model. Models (eq 1—5) correspond to MM: McConnaughey and McRoy (1979), Fry: Fry (2002), Post: Post et al. (2007), Logan: Logan et al. (2008), and Lesage: Lesage et al. (2010), respectively. Eq. 6 refers to nitrogen lipid normalization (this study).

| models | global approach scenario |  |  |  | cluster-based scenario |  |  |  | species-specific scenario |  | original coefficients scenario |  |  |  |
| --- | --- | --- | --- | --- | --- | --- | --- | --- | --- | --- | --- | --- | --- | --- |
| | mae | $\text{diff}_{\text{mae}}$ | acc. | $\text{diff}_{\text{acc}}$ | mae | $\text{diff}_{\text{mae}}$ | acc. | $\text{diff}_{\text{acc}}$ | mae | acc. | mae | $\text{diff}_{\text{mae}}$ | acc. | $\text{diff}_{\text{acc}}$ |
| MM [eq 1] | 0.39 | 0.10 | 34.6 | -11.6 | 0.31 | 0.02 | 40.9 | -5.3 | 0.29 | 46.2 | 1.50 | 1.21 | 0.9 | -45.3 |
| Fry [eq 2] | 0.38 | 0.04 | 34.2 | -5.9 | 0.35 | 0.01 | 37.6 | -2.5 | 0.34 | 40.1 | 0.38 | 0.04 | 35.4 | -4.7 |
| Post [eq 3] | 0.38 | 0.09 | 35.2 | -9.5 | 0.32 | 0.03 | 40.5 | -4.2 | 0.29 | 44.7 | 0.62 | 0.33 | 15.3 | -29.4 |
| Logan [eq 4] | 0.37 | 0.08 | 34.8 | -10.5 | 0.31 | 0.02 | 41.1 | -4.1 | 0.29 | 45.2 | 0.39 | 0.10 | 35.5 | -9.7 |
| Lesage [eq 5] | 0.43 | 0.13 | 30.2 | -13.6 | 0.36 | 0.06 | 37.1 | -6.7 | 0.30 | 43.8 | 1.49 | 1.19 | 2.3 | -41.5 |
| Nitrogen [eq 6] | 0.39 | 0.09 | 40.4 | -12.0 | 0.31 | 0.01 | 52.0 | -0.4 | 0.30 | 52.4 | - | - | - | - |

Model fit was also consistently best when δ^13^C was normalized using model coefficient values optimized with our dataset, as in the cluster-based scenario, instead of when using the original published values, like in the original coefficients scenario. The gain in model performance from optimized coefficient values was the greatest in MM and Lesage models, and the smallest in Fry and Logan models. Model coefficient values optimized for each combination of cluster and model are shown in Table 4. Coefficient D, involved in the MM and Fry models, stands as a discrimination factor between pure proteins and pure lipids and was assigned a value of 6‰ by McConnaughey and McRoy (1979) and by Fry (2002). The D value optimized with our dataset fluctuated across clusters but was generally lower than 6‰, and was lower for the MM model than the Fry model (Table 4). Coefficient values belonging to the species-specific scenario, are shown in Appendix S1 (supporting information).

MM and Logan models showed consistently the lowest mean absolute errors and the highest relative accuracy in all scenarios except in the original-coefficients scenario. Comparison of model performances within clusters further confirmed MM and Logan as the best options for lipid-normalizing δ^13^C of marine species (Table 3; Appendix S2 in Supporting information). Both models were the highest or second highest ranking models in each cluster except one (cluster A for MM and cluster G for Logan). The Lesage model showed low AIC values but all other indicators showed poor model performance (Table 3, Appendix S2 in Supporting information).

**Table 3.** Identification of the most performant normalization models, using three criteria and eight datasets (clusters A-H in cluster-based scenario): 1) CI: whether the 95 % confidence interval of the slope of a linear relationship between predicted δ^13^C_lipid-free_ and observed δ^13^C_lipid-extracted_ includes (CI = y) or not (CI = n) the target value of 1.00; 2) slope: rank based on the increasing absolute distance to target value of 1.00; 3) mae: rank based on increasing mean absolute error. Bold indicate the best and second-best ranking models. Models (eq 1—5) correspond to MM: McConnaughey and McRoy (1979), Fry: Fry (2002), Post: Post et al. (2007), Logan: Logan et al. (2008), and Lesage: Lesage et al. (2010), respectively.

| Cluster | n | MM [eq 1] | Fry [eq 2] | Post [eq 3] | Logan [eq 4] | Lesage [eq 5] |
| --- | --- | --- | --- | --- | --- | --- |
|  |  | CI ; slope ; mae | CI ; slope ; mae | CI ; slope ; mae | CI ; slope ; mae | CI ; slope ; mae |
| A | 205 | n;2;3 | **n;1;2** | **n;1;1** | **n;1;2** | n;3;4 |
| B | 420 | **y;1;1** | n;3;3 | y;2;2 | **y;2;1** | n;4;4 |
| C | 339 | **n;1;2** | **n;1;2** | n;1;3 | **n;1;2** | **n;2;1** |
| D | 413 | **n;1;2** | n;2;4 | **n;1;1** | **n;1;2** | n;3;3 |
| E | 251 | **y;2;2** | **y;1;3** | **y;2;2** | **y;2;2** | **n;3;1** |
| F | 237 | **n;1;2** | n;1;3 | **n;1;1** | **n;1;2** | n;2;2 |
| G | 67 | **y;2;1** | **y;1;1** | y;3;3 | y;3;2 | n;4;4 |
| H | 422 | **n;3;2** | **n;1;4** | **n;2;3** | **n;2;1** | n;4;5 |

**Table 4.**
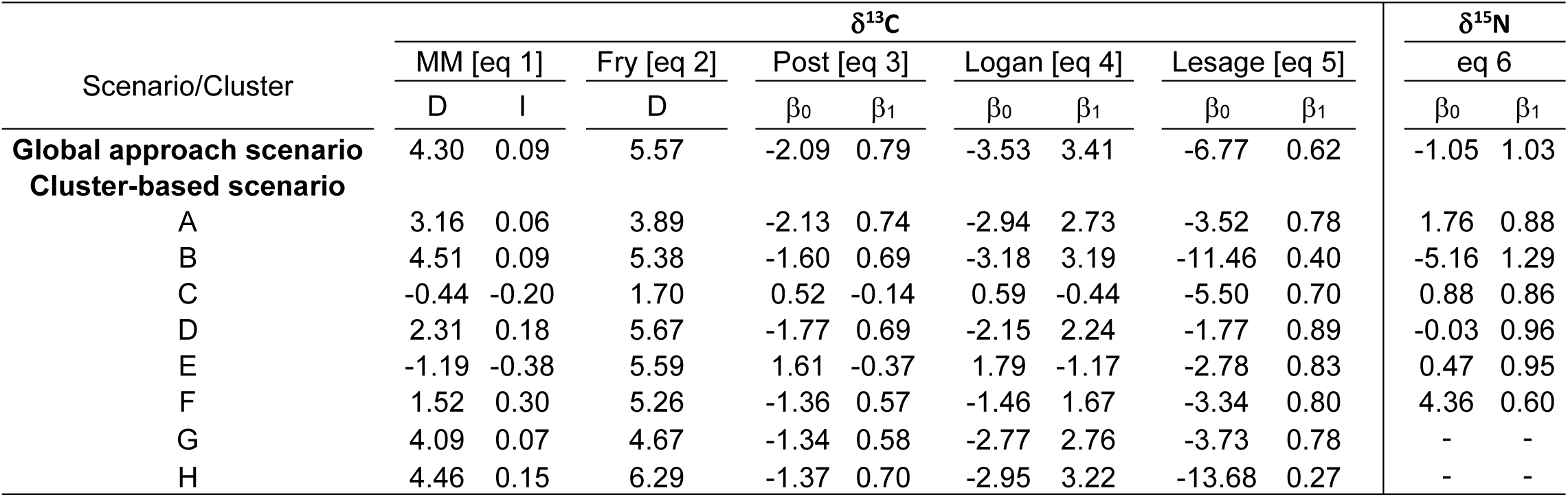
Coefficient estimates of the five lipid-normalization models for δ^13^C values, and the model retro-correcting δ^15^N_lipid-extracted_ values. Model references are as follows for carbon lipid-normalization MM : McConnaughey and McRoy (1979), Fry : Fry (2002), Post: Post et al. (2007), Logan: Logan et al. (2008), Lesage : Lesage et al. (2010). Eq. 6 refers to nitrogen retro-correction (this study).

The δ^15^N retro-correction model generally led to a slightly superior accuracy than δ^13^C lipid-normalization models but had mean absolute error values well within the range of δ^13^C lipid-normalization models in all scenarios (Table 2). In the global approach scenario, the retro-correction of δ^15^N achieved a greater r^2^ but a lower slope than lipid-normalization models (Appendix S2 in Supporting information). Differences in cluster composition between the C- and N-classifications prevented a direct comparison of model performance indicators per cluster in the cluster-based scenario. However, the retro-correction models of δ^15^N for the different clusters had globally performance indicators that were comparable to those obtained for lipid-normalization of δ^13^C (Appendix S2 in Supporting information), and in some cases, but not all, a lower mean absolute error and greater r^2^ than the model combining all species together.

## 4. DISCUSSION

This study advances the field of trophic ecology using isotopic tracers by providing a review of the performance of existing lipid-normalization models using a large dataset (i.e., over 2000 samples) of 28 marine species. Our results indicate that while there are clear benefits in exploiting taxa- or group-specific models, some models such as the one developed by Logan et al. (2008) not only outperformed several other models, but introduced little additional error. We propose a way forward for how to best deal with lipids and lipid extraction biases in isotopic studies, including a most-needed model to restore old datasets where lipid-extraction was applied systematically to all samples, to revert δ^15^N_lipid-extracted_ values back to their bulk values.

### 4.1 Effect of lipid extraction

Our study brought further evidence that lipid extraction results in an enrichment (positive discrimination) of both δ^13^C and δ^15^N values, and that in all but a handful of species, this change exceeded the measurement error (Post et al. 2007, Logan et al. 2008, Choy et al. 2016, Clark et al. 2019, Lerner & Hunt 2022). Expectedly, species with the largest discrimination in ^13^C were those with the highest C:N, which are often – but not always - the most lipid-rich samples (Fischer-Rush et al. 2021, but see Fagan et al. 2011, Yurkowski et al. 2014, Choy et al. 2016). The C:N has been used extensively in the literature to determine whether lipid-extraction or normalization was warranted or not. Generally, samples characterized by low C:N_bulk_ (<3.5) and low lipid content (<15%) were considered to be minimally affected by lipid extraction (Post et al. 2007, Logan et al. 2008, Skinner et al. 2016). Our study, however, demonstrates a wide spread of Δδ^13^C estimates at low C:N values (Fig. 1c), and argues against such an approach. Other authors also warned against not accounting for lipids at low C:N values, and recommended that the δ^13^C depletion bias in lipids be always accounted for as a precaution (Choy et al. 2016, Rioux et al. 2019, Cloyed et al. 2020, Groß et al. 2021). δ^13^C values of grey seal (*Halichoerus grypus*) were notably not affected by lipid extraction and this was also reported by Clark et al. (2019) in muscle, and only in muscle, of walrus (*Odobenus rosmarus*), another pinniped species. These authors explained this result with a low lipid content in muscle tissue. Similarly, Cloyed et al. (2020) found no effect of lipid extraction on δ^13^C in West Indian manatee (*Trichechus manatus*) muscle. The positive relationship observed between δ^13^C_bulk_ and C:N_bulk_ in cluster E, consisting only of northern shrimp (*Pandalus borealis*), was atypical compared to other clusters. However this was only a weak relationship with a high *p*-value (0.09) and low r^2^ (0.01) and a narrow range of C:N_bulk_ values. Although C:N_bulk_ was not a good predictor for δ^13^C_bulk_ in this particular species, lipid-normalization model performed adequately.

### 4.2 Model performances

The linear relationship observed between bulk and lipid-extracted δ^15^N was consistent with multiple previous studies (i.e., Logan et al. 2008, Lesage et al. 2010, Groß et al. 2021) and confirmed the possibility to retro-correct with high accuracy the δ^15^N values using the isotopic ratio measured on lipid-extracted aliquots.

Most lipid-normalization models for δ^13^C could predict δ^13^C_lipid-free_ with a satisfactory level of accuracy. The exception that stood out was the Lesage model (eq. 5), possibly because this model is specific to cetacean skin instead of muscle or whole organism, or because it omits the C:N parameter. We therefore recommend against the use of this model for estimating δ^13^C_lipid-free_ in muscle tissue, and instead suggest using the Logan or MM models as they performed generally better in most cases, despite small differences favoring the Logan model overall.

### 4.3 Model specificity

Increasing model specificity from a generalized model to a species-specific model or by using coefficients optimized for the study dataset, reduced the mean absolute error in all models and for both isotopes, a finding that is consistent with previous studies (Logan et al. 2008, Lesage et al. 2010, Cloyed et al. 2020). Our results, however, indicate that an intermediate scenario where species are grouped based on their C:N (cluster-based scenario) may still achieve high accuracy without the drawbacks and labor-intensity and costs of an approach where each species is analyzed individually, with the additional issues arising with rare species and small sample size.

Models for retro-calculating δ^15^N_bulk_ performed slightly better than the δ^13^C lipid-normalization models for the three scenarios (i.e., global approach, cluster-based, species-specific). Worth noting is the fact that model fit was higher for the cluster-based scenario than the global model (global approach scenario) in some, but not all cases. These results are important as they can help define a way forward to deal with lipid effects or lipid-extraction biases in isotopic studies depending on the specifics of each dataset.

The decision tree for how to proceed depends largely on species identity. In cases where the species exists in our dataset, isotopic analyses of lipid-extracted samples with subsequent correction of δ^15^N_lipid-free_ values using the model and coefficient most suited to the species under study (Fig. 3, Appendix S2, Table 4), would achieve the highest accuracy. An alternate approach with a slightly lower accuracy but higher labor-efficiency would be to proceed with isotopic analyses on bulk samples, with a subsequent lipid-normalization of δ^13^C_bulk_ values using again, the model (likely the Logan model) and coefficient most suited to the species under study.

In cases where the species does not exist in our dataset, the investigator’s decision could go two ways. In cases where the C:N_bulk_ would not be expected to be out of range compared to values in our study, the preferred approach would be to proceed with isotopic determination from lipid-extracted samples, with subsequent restoration of δ^15^N_bulk_ using the generalized equation, which achieves higher accuracy than lipid-normalization based on all samples combined, but with acceptance of a potential loss of accuracy for some species over a cluster-based approach. Alternatively, the investigator could proceed with isotopic determination from bulk samples. For species with a C:N_bulk_ and δ^13^C_bulk_ values within the range of the species examined in our study, investigators could attempt a correction by selecting the appropriate coefficients and cluster based on the mean C:N_bulk_ and δ^13^C_bulk_ of their species. We recommend against opting for the most phylogenetically-related species since there was no apparent phylogenetic kinship in the clusters composition. If C:N_bulk_ value of the species ends up to be out of range, the investigator should, as a last resort, use the Logan (or MM) model and associated coefficients for all species combined (global approach scenario), and accept some loss in precision.

This study provides a comprehensive and critical review of existing approaches to lipid effects in isotopic studies, and proposed a clear way forward for obtaining reliable δ^13^C and δ^15^N values without the costs from duplicate analyses. As such, it represents a major step toward the harmonization of datasets collected as part of long-term studies.

## ACKNOWLEDGEMENTS.

The authors wish to thank the many people and organizations who have participated in bringing together this vast collection of samples, especially C. Nozères, Y. Morin, K. Gavrilchuk, D. Archambault, G. Chouinard, S. Bérubé, H. Bourdages, M. Hammill, S. Mongrain, P. Joly, S. Plourde, P. Rivard, G. Cyr, R. Dubé, B. Sainte-Marie, and the field crew of the CCGS Teleost, CCGS Leim and Lampsilis, the Québec Marine Mammal Emergency Response Network, Mingan Islands Cetacean Study, J. Gauthier (Pêcheries Charlevoix), P. Lizotte (Pêcheries Lizotte), and Ministère des Forêts, de la Faune et des Parcs du Québec. Sample preparation was conducted by DFO personnel : Y. Morin, D. Gaspard, C. Potvin, K. Gavrilchuk, S. Aucoin, L. Bennour, M.-È. Chartrand-Lemieux, J. Riopel, C. Tessier-Larivière, M. Unger, P. Gagnon. Training in mixtures models was provided by Gabriel Pigeon. We also thank R. Drimmie and M. Stas, and the technical staff from the Isotope lab at University of Waterloo, and Isotope Tracer Technologies Inc. (Ontario, CAN) for the isotopic analyses. This research project was funded under the St. Lawrence Action Plan-Vision 2000, Species at Risk program and Whales Initiatives research program of Fisheries and Oceans Canada, the Green Plan of Parks Canada, and the Mingan Islands Cetacean Study. Specimens and sample collection were entirely conducted under permits delivered by the Department of Fisheries and Oceans of Canada.

## AUTHORS CONTRIBUTION

V.L. and J.C. designed the study. All authors participated in writing the final manuscript, organizing sample collection, maintenance of the dataset, and interpretation of the results. X.B., J.C., V.L. and E.R. conducted sample preparation prior to SIA. J.F.O. tested the models and analyzed the data. Approval for publication was granted by all authors.

## DATA AVAILABILITY STATEMENT

Data presented in this study is archived in Government of Canada facilities and is available upon request from Véronique Lesage.

