## supporting information 1 and 2 for "Dealing with lipid effects and lipid-extraction biases in δ^13^C and δ^15^N isotopic studies: a solution based on 28 marine invertebrate, fish and mammal species"

Appendix S1. Coefficient estimates of the five lipid-normalization models for $\delta^{13}C$ values, and the model retro-correcting $\delta^{15}N_{lipid\text{-}extracted}$ values, for each species considered individually (species-specific scenario). Model references are as follows for carbon lipid-normalization MM : McConnaughey and McRoy (1979), Fry : Fry (2002), Post: Post et al. (2007), Logan: Logan et al. (2008), Lesage : Lesage et al. (2010). Eq. 6 refers to nitrogen retro-correction (this study).

| | $\delta^{13}C$ | | | | | | | | | $\delta^{15}N$ | |
|---|---|---|---|---|---|---|---|---|---|---|---|
| Species | MM [eq 1] | | Fry [eq 2] | Post [eq 3] | | Logan [eq 4] | | Lesage [eq 5] | | eq 6 | |
| | D | I | D | $\beta_0$ | $\beta_1$ | $\beta_0$ | $\beta_1$ | $\beta_0$ | $\beta_1$ | $\beta_0$ | $\beta_1$ |
| American eel (*Anguilla rostrata*) | 3.81 | 0.08 | 4.33 | 0.13 | 0.31 | -1.41 | 1.92 | 0.66 | 0.95 | 0.84 | 0.90 |
| American plaice (*Hippoglossoides platessoides*) | 2.30 | 0.17 | 4.58 | -1.91 | 0.73 | -2.23 | 2.28 | -2.35 | 0.86 | 3.68 | 0.70 |
| Arctic cod (*Boreogadus saida*) | 4.89 | 0.09 | 8.31 | -3.46 | 1.25 | -4.55 | 4.35 | -6.15 | 0.67 | 0.83 | 0.89 |
| Arctic krill (*Thysanoessa spp.*) | 2.34 | 0.10 | 2.95 | -1.46 | 0.55 | -2.04 | 1.99 | -5.01 | 0.70 | 2.80 | 0.66 |
| Atlantic cod (*Gadus morhua*) | 1.26 | 0.28 | 4.22 | 0.60 | -0.06 | 0.62 | -0.18 | 0.94 | 1.03 | 0.28 | 0.94 |
| Atlantic herring (*Clupea harengus*) | 4.63 | 0.08 | 5.25 | -1.16 | 0.61 | -2.97 | 3.07 | -13.26 | 0.30 | 4.31 | 0.60 |
| Atlantic mackerel (*Scomber scombrus*) | 3.01 | 0.13 | 5.53 | -2.17 | 0.81 | -2.78 | 2.76 | -5.03 | 0.73 | 1.22 | 0.83 |
| Atlantic seasnail (*Liparis atlanticus*) | 3.26 | 0.12 | 9.56 | -2.49 | 0.91 | -3.09 | 3.03 | -6.59 | 0.64 | 2.24 | 0.79 |
| Atlantic tomcod (*Microgadus tomcod*) | -1.87 | -0.16 | 1.54 | 1.94 | -0.52 | 2.29 | -1.73 | -0.09 | 0.98 | 1.08 | 0.89 |
| beluga (*Delphinapterus leucas*) | 3.89 | 0.02 | 4.27 | -2.54 | 0.84 | -3.62 | 3.23 | -13.10 | 0.24 | 1.76 | 0.88 |
| capelin (*Mallotus villosus*) | 2.48 | 0.31 | 6.38 | -1.07 | 0.58 | -1.71 | 2.15 | -5.37 | 0.69 | 3.87 | 0.64 |
| eelpout (*Lycodes sp*) | 6.09 | 0.00 | 5.32 | -5.14 | 1.63 | -6.38 | 5.55 | -10.28 | 0.43 | 3.54 | 0.71 |
| Greenland halibut (*Reinhardtius hippoglossoides*) | 4.61 | 0.16 | 6.26 | -0.57 | 0.56 | -2.38 | 2.92 | -13.20 | 0.28 | 1.73 | 0.80 |
| grey seal (*Halichoerus grypus*) | 1.93 | -0.04 | 1.03 | -1.82 | 0.55 | -2.16 | 1.81 | -8.55 | 0.55 | 1.10 | 0.91 |
| longhorn sculpin (*Myoxocephalus octodecemspinosus*) | 0.82 | 0.75 | -12.64 | -0.50 | 0.36 | -0.48 | 0.96 | -5.95 | 0.65 | -0.09 | 0.94 |
| northern krill (*Meganyctiphanes norvegica*) | 6.88 | 0.04 | 6.66 | -4.52 | 1.56 | -6.30 | 5.79 | -8.94 | 0.52 | 2.23 | 0.74 |

| | | | | | | | | | | | |
|---|---|---|---|---|---|---|---|---|---|---|---|
| northern shortfin squid (*Illex illecebrosus*) | 4.08 | 0.14 | 7.66 | -2.78 | 1.07 | -3.63 | 3.66 | 0.72 | 0.99 | 7.41 | 0.39 |
| northern shrimp (*Pandalus borealis*) | -1.19 | -0.38 | 5.59 | 1.61 | -0.37 | 1.79 | -1.17 | -2.78 | 0.83 | 4.65 | 0.58 |
| rainbow smelt (*Osmerus mordax*) | -2.52 | -0.14 | 2.08 | 2.68 | -0.74 | 3.10 | -2.40 | 0.12 | 0.99 | 0.85 | 0.90 |
| redfish (*Sebastes sp*) | 3.65 | 0.08 | 5.52 | -2.76 | 0.96 | -3.51 | 3.30 | -6.04 | 0.67 | -0.80 | 1.01 |
| sand lance (*Ammodytes sp*) | -0.98 | -0.68 | 6.39 | 1.58 | -0.29 | 1.74 | -0.93 | -1.16 | 0.91 | 0.33 | 0.91 |
| shorthorn sculpin (*Myoxocephalus scorpius*) | 2.98 | 0.15 | 2.46 | -3.05 | 1.10 | -3.25 | 3.21 | -2.97 | 0.81 | 5.38 | 0.60 |
| silver hake (*Merluccius bilinearis*) | 5.06 | 0.03 | 5.42 | -3.34 | 1.13 | -4.64 | 4.20 | -12.77 | 0.33 | -1.57 | 1.06 |
| striped bass (*Morone saxatilis*) | 4.89 | 0.05 | 5.50 | -3.79 | 1.28 | -4.82 | 4.41 | 0.06 | 0.98 | 0.13 | 0.97 |
| thorny skate (*Amblyraja radiata*) | 2.87 | 0.38 | -1.88 | -3.42 | 1.51 | -3.03 | 3.68 | -8.68 | 0.51 | 0.20 | 0.97 |
| white hake (*Urophycis tenuis*) | -2.96 | -0.08 | 12.67 | 3.25 | -0.95 | 3.64 | -2.95 | -0.14 | 0.97 | -0.32 | 0.97 |
| winter flounder (*Pseudopleuronectes americanus*) | 1.01 | 0.72 | 6.57 | -0.66 | 0.43 | -0.65 | 1.20 | -8.32 | 0.53 | -5.16 | 1.29 |
| witch flounder (*Glyptocephalus cynoglossus*) | 1.26 | 0.28 | 4.42 | -1.12 | 0.47 | -1.20 | 1.36 | -3.24 | 0.80 | 4.53 | 0.66 |

Appendix S2. Model performance parameters [95% slope confidence interval] estimates from the linear relationship between predicted $\delta^{13}C_{lipid\text{-}free}$ and observed $\delta^{13}C_{lipid\text{-}extracted}$ and between predicted and observed $\delta^{15}N_{bulk}$ for the global approach and cluster-based scenarios, mae: mean absolute error. Models (eq 1—5) for carbon normalization correspond to MM: McConnaughey and McRoy (1979), Fry: Fry (2002), Post: Post et al. (2007), Logan: Logan et al. (2008), and Lesage: Lesage et al. (2010), respectively. Cluster composition differs between C and N-classifications, therefore indicators cannot be compared within clusters.

| Scenario/ Cluster | Indicator | **$\delta^{13}C$** MM [eq 1] | Fry [eq 2] | Post [eq 3] | Logan [eq 4] | Lesage [eq 5] | *n* | $C:N_{bulk}$ range | **$\delta^{15}N$** eq 6 | *n* |
|---|---|---|---|---|---|---|---|---|---|---|
| **Global approach scenario** | | | | | | | | | | |
| | slope | 0.93 [0.91—0.95] | 0.96 [0.94—0.98] | 0.99 [0.97—1.01] | 0.98 [0.96—1.00] | 0.69 [0.67—0.71] | 2354 | 2.30—8.90 | 0.89 [0.88—0.91] | 2354 |
| | mae | 0.39 | 0.38 | 0.38 | 0.37 | 0.43 | | | 0.39 | |
| | $r^2$ | 0.75 | 0.80 | 0.81 | 0.81 | 0.69 | | | 0.89 | |
| | aic | 3804.0 | 3307.3 | 3345.9 | 3291.4 | 3120.3 | | | - | |
| **Cluster-based scenario** | | | | | | | | | | |
| A | slope | 0.90 [0.84—0.97] | 0.91 [0.84—0.98] | 0.91 [0.84—0.97] | 0.91 [0.84—0.97] | 0.71 [0.64—0.77] | 205 | 2.73—5.70 | 0.80 [0.72—0.89] | 92 |
| | mae | 0.34 | 0.33 | 0.32 | 0.33 | 0.36 | | | 0.36 | |
| | $r^2$ | 0.78 | 0.79 | 0.79 | 0.78 | 0.71 | | | 0.80 | |
| | aic | 237 | 228.8 | 222.3 | 228.7 | 211.7 | | | - | |
| B | slope | 0.95 [0.89—1.01] | 0.92 [0.86—0.99] | 0.94 [0.87—1.00] | 0.94 [0.88—1.00] | 0.35 [0.30—0.39] | 420 | 2.76—8.67 | 0.83 [0.72—0.95] | 42 |
| | mae | 0.32 | 0.38 | 0.35 | 0.32 | 0.43 | | | 0.60 | |
| | $r^2$ | 0.70 | 0.64 | 0.67 | 0.69 | 0.35 | | | 0.84 | |
| | aic | 490.3 | 579.7 | 544.1 | 498.0 | 268.7 | | | - | |
| C | slope | 0.72 [0.65—0.80] | 0.72 [0.64—0.79] | 0.72 [0.64—0.80] | 0.72 [0.65—0.80] | 0.49 [0.44—0.55] | 339 | 2.30—3.85 | 0.90 [0.86—0.93] | 289 |
| | mae | 0.32 | 0.32 | 0.33 | 0.32 | 0.31 | | | 0.36 | |
| | $r^2$ | 0.50 | 0.51 | 0.50 | 0.50 | 0.49 | | | 0.90 | |

| | | | | | | | | | | |
|---|---|---|---|---|---|---|---|---|---|---|
| | aic | 360.9 | 349.3 | 360.4 | 360.7 | 116.4 | | | - | |
| D | slope | 0.80 [0.76—0.84] | 0.78 [0.74—0.83] | 0.80 [0.76—0.84] | 0.80 [0.76—0.84] | 0.73 [0.69—0.78] | 413 | 2.59—4.24 | 0.90 [0.89—0.92] | 933 |
| | mae | 0.26 | 0.31 | 0.25 | 0.26 | 0.28 | | | 0.32 | |
| | $r^2$ | 0.77 | 0.74 | 0.78 | 0.78 | 0.73 | | | 0.90 | |
| | aic | 216.1 | 267.3 | 197.9 | 206.3 | 232.6 | | | - | |
| E | slope | 0.98 [0.92—1.04] | 1.00 [0.94—1.07] | 0.98 [0.92—1.04] | 0.98 [0.92—1.04] | 0.81 [0.76—0.86] | 251 | 2.76—3.85 | 0.92 [0.89—0.95] | 308 |
| | mae | 0.28 | 0.31 | 0.28 | 0.28 | 0.26 | | | 0.20 | |
| | $r^2$ | 0.82 | 0.80 | 0.82 | 0.82 | 0.81 | | | 0.92 | |
| | aic | 180.7 | 217.3 | 180.3 | 180.5 | 90.3 | | | | |
| F | slope | 0.90 [0.82—0.97] | 0.90 [0.82—0.97] | 0.90 [0.83—0.97] | 0.90 [0.83—0.97] | 0.72 [0.66—0.77] | 237 | 2.68—4.64 | 0.48 [0.45—0.52] | 690 |
| | mae | 0.29 | 0.38 | 0.28 | 0.29 | 0.29 | | | 0.30 | |
| | $r^2$ | 0.73 | 0.69 | 0.73 | 0.73 | 0.72 | | | 0.49 | |
| | aic | 263.3 | 304.0 | 255.1 | 259.5 | 170.0 | | | - | |
| G | slope | 1.03 [0.96—1.10] | 1.02 [0.95—1.09] | 1.04 [0.97—1.11] | 1.04 [0.97—1.11] | 0.91 [0.84—0.98] | 67 | 3.16—7.33 | | |
| | mae | 0.48 | 0.48 | 0.52 | 0.49 | 0.53 | | | | |
| | $r^2$ | 0.94 | 0.93 | 0.93 | 0.93 | 0.91 | | | | |
| | aic | 124.0 | 127.7 | 131.3 | 127.1 | 128.7 | | | | |
| H | slope | 0.64 [0.58—0.69] | 0.74 [0.66—0.81] | 0.66 [0.60—0.73] | 0.66 [0.60—0.72] | 0.10 [0.07—0.13] | 422 | 2.96—8.90 | | |
| | mae | 0.35 | 0.41 | 0.36 | 0.34 | 0.49 | | | | |
| | $r^2$ | 0.52 | 0.47 | 0.51 | 0.54 | 0.10 | | | | |
| | aic | 376.4 | 593.1 | 438.4 | 385.6 | -209.6 | | | | |

14
